# Near-threshold somatosensory perception revisited: pre- and post-stimulus neural processes during electrical and naturalistic mechanical stimulation

**DOI:** 10.64898/2026.11.14.765324

**Authors:** R. Großmann, S. Woo, T. R. Knösche, A. Villringer

## Abstract

Near-threshold electrical stimulation is widely used to study the post-stimulus neural correlates of conscious somatosensory perception and to examine how ongoing brain activity relates to stimulus detection. However, the precise relationship between pre-stimulus alpha activity and conscious somatosensory detection still remains debated, and a previous study suggests that a more natural mechanical somatosensory stimulation may evoke different neural responses. In a within-subject study of 40 healthy volunteers, we compared early event-related EEG responses to near-threshold mechanical and electrical stimuli and examined how pre-stimulus alpha amplitude and phase coherence related to stimulus detection. Both stimulation modalities elicited robust EEG correlates of stimulus detection likely corresponding to the N80 and N140 components. Detection related EEG responses were localised in the right (contralateral) precentral and postcentral gyri, with the left ipsilateral hemisphere contributing shortly afterwards. Detection was associated with lower pre-stimulus alpha amplitude and greater alpha phase coherence. We found no evidence that the early detection-related EEG responses or the associations with pre-stimulus alpha differed between stimulation modalities.

## Introduction

The identification of neural correlates of somatosensory perception, which reflect the minimal processes giving rise to a conscious experience of the stimulus [1], is a long-standing central interest in neuro-science. Near-threshold paradigms provide a valuable approach to investigating these relationships by enabling comparisons between perceived and unperceived stimuli of intensities near the perceptual threshold. Those paradigms require overt responses (e.g., button press, saccades) from subjects to categorize a somatosensory stimulus as perceived. Previous studies using near-threshold paradigms with electrical stimulation consistently identified a mid latency component (around 140 ms post stimulus) as a promising correlate for conscious perception in humans [2–4], while ruling out the later component P300, assigning it to task relevance [5] and earlier components which seem to reflect physical stimulus properties like intensity [5, 6].

More recently, an invasive stereotactic electroencephalography (sEEG) study identified an even earlier neural correlate of near-threshold stimulus perception, with the first significant association between neural activity and perceptual report emerging at approximately 100 ms [7]. The authors attributed this correlate to sustained activity in the parietal operculum, a region encompassing the secondary somatosensory cortex (S2). These findings are not restricted to near-threshold paradigms. Using a masking paradigm, Schubert et al. [8] likewise found comparable P50 and N80 amplitudes for detected and undetected stimuli, whereas later mid-latency somatosensory evoked potentials, including the P100 and N140, were enhanced for consciously perceived stimuli. Until now, almost all studies applying near-threshold somatosensory stimulation focused on electrical stimulation either directly at the median nerve or at the finger of the participants. Only very few studies used the more naturalistic mechanical modality for near-threshold stimulation via a piezoelectric or vibrotactile device [4, 9–11]. The first study comparing mechanical sub- and supra-threshold stimulation found the ERP components P50, P100, N190 for supra-threshold, but not for sub-threshold stimuli [9]. Not accounting for stimulus intensity means that they could not test whether the absence of each component was due to perceptual processes or stimulus intensity. Additionally, they did not directly compare to electrical stimulation, which limits their interpretation. Jones et al. [10] applied a tactile stimulus of 10 ms at the perceptual threshold using a piezoelectric device and observed that the ERP amplitude predicts somatosensory perception starting at 70 ms post-stimulus. In line with previous studies they also liked a perceptual correlate at 135 ms to repeated output from the granular layer in the primary somatosensory cortex (S1). Gusso et al. [11], using vibrotactile stimuli, found that the N80 and N140 components were associated with perceived stimuli, whereas unperceived trials were associated with the P100, with both components also showing associations with the P300. The earliest component associated with perception was the N80, occurring at approximately 75 ms. Recent findings show that the neural correlates of somatosensory perception might differ between mechanical and electrical modalities, particularly in early-to-mid latency processing [4]. The study compared electrical and mechanical near-threshold stimulation and reported that in the mechanical modality early activity at 50-60 ms was best explained by stimulus detection, while for electrical stimulation, stimulus intensity best explained that activity. Förster and colleagues argue that these differences in processing may stem from receptor specificity since electrical stimulation directly activates peripheral nerve fibers and can recruit different classes of sensory afferents depending on stimulation parameters, which may therefore contribute to an early evoked component even for subthreshold stimuli [4]. On the other hand, mechanical stimulation primarily engages cutaneous mechanoreceptors, including Meissner’s corpuscles and Merkel cell complexes in the superficial skin, as well as deeper Pacini and Ruffini corpuscles [12, 13], which might not lead to sufficient activation to elicit a P50 component. Taken together, these findings suggest that the neural processing underlying conscious perception may differ between mechanical and electrical stimulation.

However, conscious perception is not determined solely by post-stimulus processing. Pre-stimulus neural activity modulates cortical excitability and sets the stage for subsequent somatosensory perception [14, 15]. Until now, the effects of pre-stimulus alpha amplitude and phase on mechanical somatosensory detection, including a direct comparison with electrical stimulation, have not been investigated. The mu rhythm in the somatosensory system (i.e., Rolandic alpha) was found to be modulated by attention [16] and its effect on perception has been extensively studied. Its amplitude was found to correlate with stimulus detection in an inverted U-shape manner (medium amplitude leading to highest detection rates) by some studies [3, 17, 18], while others found reduced pre-stimulus alpha power for perceived compared to unperceived trials [19–21]. Moreover, medium and high amplitude Rolandic alpha in the pre-stimulus interval induced a reduction in the P50 component, when the stimulated finger was unattended, and an increase of the P50 when it was attended [3].

Besides the alpha oscillation amplitude, the alpha phase also matters for successful stimulus detection. It was found that the phase of higher amplitude alpha oscillations is predictive for the perception of stimuli [18], and that early post-stimulus phase locking predicted stimulus detection [19]. In visual detection paradigms, the effect of alpha phase was studied early on, revealing that the phase of the alpha wave directly influences perceptual gating of the stimulus [22]. Research suggests that the brain uses alpha oscillations to carefully time stimulus processing through silencing unwanted and selecting wanted activity [23]. Ongoing pre-stimulus alpha phase alignment enhances stimulus processing, by increasing the amplitude of the P1 evoked component [24], typically peaking between 100 and 150 ms post-stimulus, preceding the visual awareness negativity (VAN) [25, 26]. Fellinger and colleagues suggest that alpha phase tuning prior to stimulation enables cognitive processing of the incoming information [24]. Gruber and colleagues [27] tested if participants react faster to a stimulus when the alpha phase is aligned to the P1 peak evoked by a visual stimulus, and indeed the reaction times of responding to the target stimulus decreased in comparison to the misaligned trials. While most of these findings on temporal prediction through alpha come from the visual domain, it is likely that similar mechanisms exist in the somatosensory domain with respect to the mu rhythm. So far, pre-stimulus preparatory activity has only been examined in electrical stimulation. Rolandic alpha phase alignment and its role in temporal tuning remain unexplored in the somatosensory domain.

In this study we investigate pre-stimulus and post-stimulus processes related to conscious somatosensory perception in a within-subject design applying both electrical and mechanical stimulation. Specifically, we investigate 1) post-stimulus neural correlates of conscious perception, 2) the effect of pre-stimulus alpha amplitude on perception, and 3) the effect of pre-stimulus alpha phase on perception.

## Methods

### Participants and recording setup

We recruited 40 healthy subjects (25 females), aged between 18 and 40 years, to take part in our experiment. We conducted electroencephalography (EEG) recordings (system NeurOne Tesla, Bittium, Oulu, Finland) and measured at a sampling rate of 5000 Hz. The EEG caps (EasyCap, Herrsching, Germany) comprised 62 electrodes following the classical 10-10 montage and using POz as the ground and FCz as the reference. The subjects had no history of neurological or psychiatric conditions, brain injuries, or neurosurgical interventions. Their handedness was tested using the Edinburgh-Handedness-Questionnaire [28] and only right-handed participants were included. Each participant took part in two measurement sessions, one with electrical and one with mechanical stimulation. Two participants could not attend the mechanical session and one participant did not attend the electrical session (resulting in 38 mechanical and 39 electrical sessions). Before analysis we excluded experimental blocks based on bad fits of their psychometric curves (Fig. 1E). Blocks with a goodness of fit of *r*^2^ *<* 0.6 were excluded. One mechanical session was removed entirely due to bad psychometric curve fits in every block. Furthermore, we excluded one mechanical session due to technical issues and another one due to extensive movement of the mechanically stimulated finger. Eventually, we included 35 mechanical sessions and 39 electrical sessions in the broad band data analysis. Stricter exclusion criteria apply for the alpha oscillation analysis and source reconstruction (see the respective Methods sections). The study was approved by the local ethics committee at the medical faculty of the University of Leipzig.

**Figure 1:**
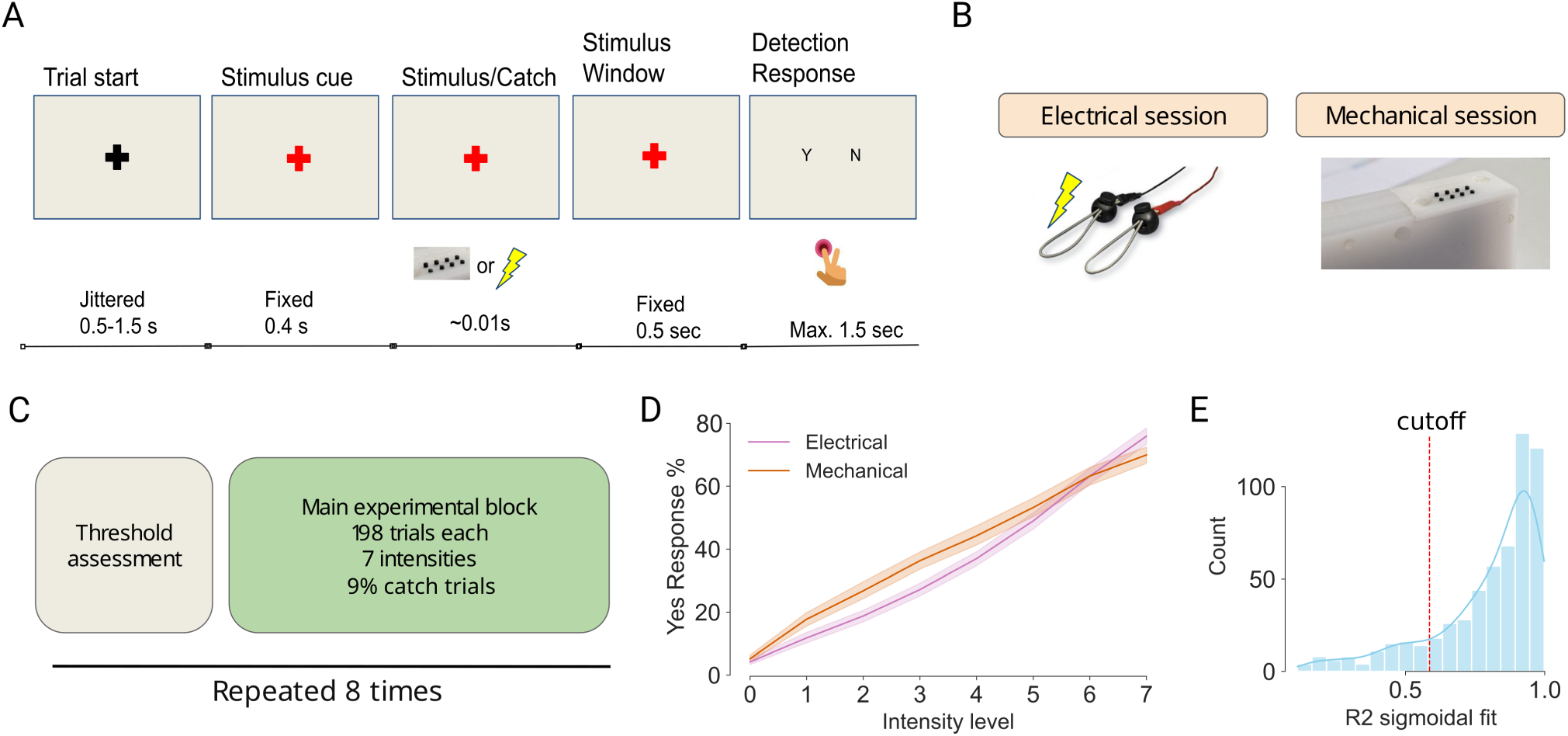
A) Trial sequence of the experiment applying either electrical (B left) or mechanical (B right) stimulation depending on the experimental session. C) Structure of one experimental session, entailing 8 experimental blocks. D) Averaged behavioral responses by intensity and stimulation modality. E) Sigmoidal fits (r-squared) computed from response curves by subject and experimental block. The red dashed line indicates the preregistered cut-off value of 0.6.

### Study design

The study follows a within-subject design, with the factors **stimulus modality** and **intensity**. Besides EEG, we recorded ECG, respiration activity using a respiration belt, and behavioral responses reflecting stimulus detection via button press with the right hand. For each participant, the two sessions were at least one day apart. After every two participants, the order of the sessions was reversed (either mechanical or electrical session first). Each session consisted of 8 experimental blocks. Each block started with a threshold assessment (4-5 min), followed by the main experimental block (9-10 min) in which all possible intensities were presented. The threshold assessment was based on the PSI-marginal adaptive algorithm [29] followed by 10 trials testing the estimated near-threshold. Manual adjustment of the threshold based on the response distribution and test result was occasionally necessary. Each main experimental block included 198 trials, with 7 different intensities, assessed to correspond to 5%, 20%, 35%, 50%, 65%, 80%, 95% yes-responses, and 18 trials without any stimulation (catch trials). The trials were presented in pseudorandomized order. For every other participant, the assignment of response buttons was swapped. Both the threshold assessment and the experimental block followed the same trial sequence. It started with a jittered inter-trial interval of 0.5-1.5 s showing a black fixation cross, followed by a red fixation cross (stimulus cue). After a fixed time window, the stimulus was delivered (for more details see next paragraph). The response window started 0.5 s post-stimulus onset and lasted for 1.5 seconds. Participants reported perception of the stimulus by pressing the corresponding button with either the right index or middle finger.

### Stimulation method

The electrical stimulation was executed using electrodes that are controlled via a medically certified constant current generator (Digitimer DS5). The mechanical stimulation was executed via small plastic pins, stimulating the finger from below, controlled by a piezoelectric stimulator (by QuaeroSys). The mechanical stimulation lasted maximally 10 ms, whereas the electrical stimulus lasted 0.2 ms. Due to a technical issue, the visual preparation cue in the mechanical modality was presented 36 ms earlier than in the electrical condition. Therefore, the cue-stimulus interval was 436 ms long in the mechanical condition, and 400 ms in the electrical condition. Participants wore earplugs and white noise was played during all experimental blocks for both modalities due to the sound of the piezoelectric stimulation. We asked participants if they could hear the piezostimulation device and increased the volume of the white noise until they confirmed that they could no longer hear it. All stimulations were applied on the left index finger.

### Data pre-processing and analysis

#### EEG pre-processing

We band-pass filtered the data (0.1-100 Hz for raw data, 1-40 Hz for epoched data), downsampled to 500Hz, applied detrending, and re-referenced to the average. Bad channels were first identified based on power spectral density (PSD) and subsequently using the RANSAC algorithm [30]. Data were epoched around stimulus onset from -900 ms to 300 ms with respect to stimulus onset, while ECG and respiration signals were filtered separately. Independent component analysis (ICA) was used to remove eye movement, cardiac, muscle, and line noise artifacts. The ICA was fitted (infomax algorithm, number of components set to the number explaining 99% of the variance) separately for each experimental block on the epoched data after band-pass filtering at 1-40 Hz. Components were marked for removal if they correlated with the vertical EOG or ECG channel, or if the ICLabel classifier [31] assigned them to any class other than brain or other. The correction was applied to the same 1-40 Hz filtered epochs, which were then used for all subsequent analyses. Finally, bad epochs were rejected using amplitude thresholds of 90 *µ*V for EEG channels and 250 *µ*V for the EOG channel. Baseline correction was applied for broad band analyses using a 100 ms long baseline starting at 550 ms pre-stimulus. We fitted sigmoidal curves to the behavioral data per block to assess their quality. Based on results from pilot experiments and preregistered, we excluded blocks with r-squared*<*0.6. Furthermore, we excluded trials applying the lowest and highest intensity level from the analysis,because there were only 15 trials of those intensities per experimental block (and due to the extreme intensity even fewer trials would be left per detection response). Because the visual cue elicits an event-related potential which dominates neural activity in the pre-stimulus interval, we used catch trials to subtract the visual ERP from each trial. Due to the low intensity of the stimuli, no omission response post-stimulus was expected which could have leaked into the pre-stimulus interval. Catch trials were not used in the analyses other than for this correction. For the pre-stimulus alpha analysis, it was important that the visual signal was removed well. This required a good signal-to-noise ratio (SNR) in the pre-stimulus window, because the visual evoked response was subtracted as a per subject average signal (see Fig. 8). We obtained the SNR by computing the root-mean-square (RMS) of the baseline signal (100 ms pre cue) and the pre-stimulus signal (400 ms post cue) of the visual ERP elicited by the visual stimulus cue for both modalities. We excluded the mechanical sessions of three subjects and the electrical sessions of three subjects (of which one subject failed the criterion for both sessions) prior to the pre-stimulus alpha analysis because their SNR was below 1.5 at visual electrode O2. The SNR did not differ significantly between modalities.

#### Statistical analysis of broad band data

The goal was to investigate the effect of stimulus modality, detection response, stimulus intensity and their interaction on the neural activity during pre- and post-stimulus intervals. Following previous studies [4, 6], we first selected somatosensory channels CP4 and C6. We performed temporal cluster-based analysis by fitting mixed effects models. The fixed effect factors were *modality*, *detection response*, and *stimulus intensity* (centred). The random-effects structure comprised a by-subject random intercept and a by-subject random slope for (centred) experimental block, to account for between-subject differences in overall amplitude and for changes across blocks such as fatigue or variability in threshold assessment. Additionally, we tested the mechanical stimulation separately to investigate if the weight of the finger lying on the mechanical stimulation device had an effect on the ERP amplitude (by introducing *finger force* as extra effect).

In addition to the somatosensory electrodes we were interested in effects across the whole brain. Especially preparation for the stimulus might go beyond activation of contralateral somatosensory cortex. To correct for multiple comparisons arising from testing many electrode locations, we performed a spatio-temporal cluster-based permutation test. We used 2000 permutations over the interval -550 to 300 ms relative to stimulus onset, with clusters formed across the channel-time adjacency and thresholded two-sided at *p* < .05. The model comprised 40 participants contributing 74 sessions (35 mechanical, 39 electrical). The factors intensity level and detection are highly interdependent, since increasing intensity levels by design lead to increased probability of detection. To investigate the collinearity of the two factors in predicting the ERP amplitude, we computed the variance inflation of all factors in the model. At electrode C6, in the time window 140-150ms the variance inflation was below 1.2 for all of the factors (intensity level: 1.0989, detection response: 1.102245, modality: 1.003261) indicating low shared variance of all the factors. The correlation between intensity level and response was 0.3. For both cluster-based permutation algorithms we relied on the Unfold [32] and ClusterDepth [33] toolboxes. For the full channel spatio-temporal analysis we modified the cluster evaluation to assessing the cluster mass instead of the depth, which is only suitable for one dimensional (here temporal) clusters.

#### Source reconstruction of the cluster-permutation *z* maps

To visualise the cortical distribution of sensor-level effects, the group-level *z*-value topographies associated with significant clusters were projected into source space using individual minimum-norm inverse operators. Only model coefficients that carried at least one significant cluster were reconstructed. Source reconstruction required an individual anatomy, so only participants with a T1 scan were included, leaving 57 sessions (28 mechanical, 29 electrical) from 33 participants, 24 of whom contributed both sessions. For each session, an individual forward solution was computed using a three-layer boundary element model (BEM) derived from the FreeSurfer segmentation [34–36]. We defined the cortical source space using an ico-5 mesh (subdividing an icosahedron five times and projecting its vertices onto the cortical surface), co-registered the session-specific digitized EEG electrode positions (using the Polhemus Patriot tracker system) to the T1 MRI image, and constrained the source vertex orientations to be normal to the cortical surface. The source-space resolution comprised 10,242 candidate source locations per hemisphere. A noise covariance matrix was estimated from the pre-cue baseline window (-550 to -450 ms relative to stimulus onset) of the cleaned epochs of that session and regularised. The forward solution and noise covariance matrix were then used to construct a fixed-orientation minimum-norm estimate (MNE) inverse operator using the MNE-Python package [37]. No depth weighting was applied because we expect mainly superficial sources to contribute to the effects (specifically in the somatosensory, frontal and parietal cortex) [38–41]. The inverse operator was applied to the channel-by-time matrix of *z* values, with the regularisation parameter set to *λ*^2^ = 1*/*9. The resulting source estimate was morphed to the fsaverage surface (ico-5, identical to the resolution of individual source space) and averaged across the 57 sessions, yielding one group source estimate per coefficient. Because the sign of a fixed-orientation minimum-norm estimate follows the cortical surface normal and therefore flips between opposing banks of a sulcus, the maps are displayed as absolute source amplitude in arbitrary units, on a single scale shared across coefficients and latencies. To describe where the strongest sources lie, we selected at each latency of interest the 1% of vertices with the largest absolute source amplitude (205 of 20,484 fsaverage vertices, both hemispheres pooled) and report the share of these vertices in each hemisphere (the right hemisphere is contralateral to the stimulated hand) and in each region of the Desikan-Killiany parcellation, separately for the left and the right hemisphere.

#### Frequency band specific analysis

To investigate pre-stimulus activity in the alpha frequency band we first cropped the broad band epochs to -800 ms to 0 ms with respect to stimulus onset, so that no post-stimulus data entered the filter. The cropped epoch was mirror padded on both sides with one full epoch length, and filtered to the subject-individual alpha frequency peak (within 8 to 14 Hz) at electrode CP4 (estimated via Welch PSD over a window from -800 to -100 ms) *±* 2 Hz using a zero-phase FIR filter. The padding was removed afterwards. A surface Laplacian (current source density) transform was applied prior to all channel-level analyses to improve isolation of local activity. We computed the analytic signal via the Hilbert transform (applying mirror padding of three 8 Hz cycles), yielding trial-wise instantaneous envelope and phase. For further analysis, the alpha band trace was used only until 100 ms pre-stimulus to prevent keeping effects of potential leakage from filtering. The envelope analyses were conducted over the pre-stimulus window (-400 to -100 ms) at the contralateral somatosensory electrodes CP4 and C4 and, as a control site outside somatosensory cortex, the occipital electrode O2. Pre-stimulus alpha envelope was compared between detected and undetected trials using paired t-tests on the subject-wise means. To test for a non-linear (inverted-U) relationship between alpha amplitude and detection, trial-wise mean envelope amplitude was sorted into 10 equiprobable bins per subject using quantile binning; a quadratic polynomial was fitted to each subject’s bin-wise detection rate, and the quadratic coefficient was tested against zero using a one-sided t-test (H_1_: coefficient *<* 0). To compare the modalities directly, a linear mixed model, envelope *∼* response *×* modality + (1 *|* subject), with centred predictors was fitted per channel to the subject-level mean envelope; the p-value of each effect was FDR-corrected across the three channels. Phase locking was quantified as debiased squared intertrial phase coherence (dITPC). We chose to look at electrode CP4 based on [3] and CPz because it showed clear preparation signals during the pre-stimulus window. At every time point within a window of -750 to -100 ms, the biased ITPC is the length of the mean resultant vector of the unit-normalised complex analytic signal across trials, computed separately for perceived and unperceived trials. The squared resultant length is biased upwards when few trials are available, which would inflate the ITPC of the condition with fewer trials. We therefore corrected the computed squared length for the number of trials *N* of the condition (see equation 1). The dITPC can be negative after the upwards bias was removed by the correction.

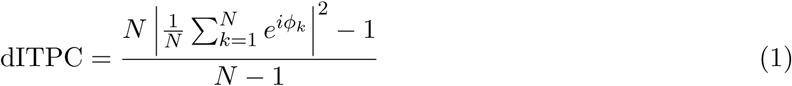

Even though the ERP evoked by the visual cue was subtracted using catch trials, there might still be a residual time-locked evoked response, which by itself raises the phase coherence across trials, so a dITPC difference between perceived and unperceived trials could reflect the ERP instead of the ongoing alpha phase. We therefore removed the ERP contribution from the dITPC time course before testing. The ERP was computed from the same epochs without the band-pass (surface Laplacian, channel selection, pre-cue baseline -550 to -450 ms) as the trial average per response, and its absolute value *|*ERP(*t*)*|* served as regressor. Per subject, modality and channel, one ordinary least squares fit dITPC(*t*) = *a* + *b |*ERP(*t*)*|* was computed over the time points of perceived and unperceived trials stacked, so that both responses share the slope *b*. The cleaned time course is

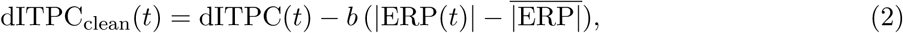

which removes the ERP term but keeps the dITPC level. All dITPC statistics below were computed on the cleaned dITPC.

Mean dITPC in the post-cue window (from cue-onset until 300 ms after cue-onset) was compared between perceived and unperceived trials using paired t-tests. To test whether the difference emerged only after the cue, the Yes-No dITPC difference in that post-cue window was additionally compared against the same difference in a pre-cue window (300 ms before cue-onset). To estimate the phase specificity independent of stimulus perception, we conducted a Rayleigh test within each modality on the phase at -100 ms, averaged within each subject. Additionally, we conducted a Rayleigh test within-subject to estimate their phase specificity across trials. To compare the post-cue dITPC effect between modalities, we fitted a linear mixed model per channel, dITPC *∼* response *×* modality + (1 *|* subject), on the subject-level post-cue mean dITPC, with response (perceived = +0.5) and modality (mechanical = +0.5) centred, so that the main effects are averaged over the other factor. The p-value of each effect was FDR-corrected across the two channels. Because phase measures depend on the exact timing between cue and stimulus, the electrical sessions of two subjects, in which the cue-stimulus interval was consistently shifted by about 8 ms relative to all other electrical sessions due to a technical issue, were additionally excluded from the dITPC analysis, leaving 34 electrical and 32 mechanical sessions. All envelope and dITPC tests were corrected for multiple comparisons across the channels tested in that analysis, within each modality, using the Benjamini-Hochberg false discovery rate (FDR) procedure. For the topographic maps of the Yes-No envelope difference, the difference was averaged across subjects first and the peak channel and latency were then identified within -400 to -100 ms.

#### Analysing the relationship between alpha effects, the P50 and the negative preparation component

To relate the pre-stimulus alpha measures to early stimulus processing we quantified the P50 at the contralateral somatosensory electrode CP4 on the broad band epochs. The epochs were baseline corrected (-550 to -450 ms). The P50 amplitude of each single trial was defined as the mean over a fixed time window, 34-56 ms for electrical and 44-66 ms for mechanical stimulation (the P50 was found to peak later in response to mechanical stimulation). As before, the alpha envelope trace was cut at -100 ms to prevent keeping effects of potential leakage from filtering in the analysis. The effect of alpha amplitude on the P50 was assessed by sorting the trial-wise pre-stimulus alpha envelope at CP4 into eight equiprobable bins within each session, so that between-subject differences in overall alpha amplitude cannot drive the binning. Bin-wise mean P50 values were averaged within subjects first and only then across subjects. As for the dITPC analysis, the electrical sessions of two subjects were excluded from these P50 analyses because of the shifted cue-stimulus interval and pre-stimulus SNR criteria, leaving 34 electrical and 32 mechanical sessions. A straight line was fitted per session across the eight bins and the resulting slopes were tested against zero with a two-sided one-sample t-test. To visualize the relationship between the alpha phase and the P50 amplitude, the instantaneous alpha phase at CP4 was read out at 100 ms before stimulation and sorted into 16 bins of equal angular width spanning the full circle. For each phase bin the P50 amplitude was computed. Within-subject averaging was applied for the phase circular mean and P50 arithmetic mean. We further compared the topographies of the pre-stimulus alpha (trial-averaged Hilbert envelope at 100 ms before stimulation) and P50 (at 42 ms and 62 ms respectively) of electrical and mechanical stimulation to test if they are likely to share their neural generator. A Laplacian was applied to both signals before computing the averages. We normalized each map by centering it, then dividing it by the root mean square of the centered map. We then applied the TANOVA method to statistically test differences in scalp topography [42]. In a second comparison, we computed per session the perceived minus unperceived difference map of the alpha envelope (-100 ms) and of the P50 (42 ms and 62 ms), normalized each map by its global field power, and tested with the same TANOVA (5000 sign-flip permutations) whether the two difference topographies differ in shape.

Finally, we quantified the slow negative preparation potential starting shortly before stimulus onset and tested its relationship with the dITPC. The negative potential peaked centrally and was therefore measured at the central electrode CPz, after low-pass filtering the broad band epochs at 6 Hz with a zero-phase FIR filter and using the same pre-cue baseline. We took the most negative sample of the trial average within -200 to 100 ms separately for perceived and unperceived trials. The dITPC was summarised per session and detection response as the maximum of the dITPC time course at CP4 within the post-cue window (-400 to -100 ms). We then correlated the dITPC with the amplitude of the negative preparation component across sessions using Pearson correlations, separately for each modality and detection response, and FDR-corrected the resulting four tests.

#### Source reconstruction of pre-stimulus alpha

To unravel the origin of alpha pre-stimulus activity in mechanical and electrical somatosensory perception, source reconstruction was performed on the same narrow band filtered epochs that were used for the channel specific pre-stimulus alpha analysis (but before the surface Laplacian transform) with the subject-specific alpha peak frequency. As for the source reconstruction of the *z*-maps, only participants with an individual anatomy were included, and sessions failing the SNR criterion or with the unintended trigger delay were excluded, leaving 52 sessions (26 electrical, 26 mechanical), 20 of whom contributed both sessions. The forward and inverse modelling approach was the same as in the previously described source reconstruction of the *z*-maps. The noise covariance matrices were estimated from the pre-stimulus baseline window (-800 ms to -450 ms with respect to stimulus on-set, long enough to capture oscillations in alpha frequency). The MNE inverse operator was applied to individual epochs, yielding trial-wise source time courses across the cortical surface, necessary for computing the dITPC. Prior to the Hilbert transform, each source time series was mirror-padded by three cycles at the lower alpha bound (8 Hz; *≈*375 ms) to suppress edge artifacts. Per source vertex we computed the Hilbert transform and obtained the envelope and the phase per trial. The envelope was averaged across trials per subject and the dITPC was computed as described in equation 1. Both measures were computed separately for perceived (“Yes”) and unperceived (“No”) trials. Individual source estimates were morphed to the fsaverage common surface and averaged across subjects. Yes-No difference maps were computed for both envelope and dITPC. Cluster-based permutation tests (2000 permutations, corrected at *p* < .05) were performed on the difference maps for each modality (over a time window of -750 ms to -100 ms relative to stimulus onset) to obtain only statistically relevant clusters.

## Results

### Behavioural Results

The experiment was designed such that the delivered intensities are tuned to the individual psychometric curve. Behavioral results confirmed the expected increase of detection upon an increase of stimulus intensity (Fig. 1D). Detection response (perceived versus unperceived) was modelled using mixed-effects logistic regression with subject-level random intercepts and block as a random slope. Across both modalities, as intended by the experiment design, stimulus intensity was the dominant predictor of detection (*β* = 0.52, SE = 0.009, z = 57.97, p *< .*001). Mechanical stimulation showed an overall higher detection rate than electrical stimulation (*β* = 0.28, SE = 0.018, z = 15.27, p *< .*001), but was still clearly at near-threshold intensity. A significant modality-by-intensity interaction (*β* = *−*0.10, SE = 0.013, z = *−*8.08, p *< .*001) indicated that the psychometric slope was steeper for electrical than for mechanical stimulation. We fitted a modality specific model to include the weight of the finger resting on the piezostimulator as an additional predictor. A higher weight of the finger might facilitate perception, as the finger is closer to the stimulation pin. This assumption was confirmed by our statistical analysis. Force quantiles explained additional variance in detection beyond the intensity level (*β* = 0.10, SE = 0.018, z = 5.61, p *< .*001), with higher force leading to increased perception.

### Broad band single channel analysis

Based on previous literature we chose to investigate the ERP traces of the electrodes CP4 and C6, which are placed above the somatosensory cortices. For each channel we fitted the same joint linear mixed model as in the whole-scalp analysis at every time point and tested the coefficients with a temporal cluster-mass permutation test (two-sided, 2000 permutations, cluster-forming threshold *z* = 1.96, corrected at *p* < .05). Significant time windows are marked below the ERP traces in Fig. 2 A and B. Table 1 lists all significant clusters at CP4 and C6 for every model coefficient, with their statistics. At both channels, detection response was associated with amplitude differences before the stimulus and with a strong post-stimulus negativity for perceived trials with two peaks in that window (often referred to as the N80 and the N140 component), which was strongest at C6. The modality effect differed at both channels before and after the stimulus. Stimulus intensity yielded clusters only at CP4, one of them covering the latency of the P50 (see Fig. 2C for the intensity effect). Modality and detection response interacted before the stimulus at CP4 and late in the epoch at C6. Intensity level and detection interacted at C6 in a mid latency time window during the N140 peak.

**Figure 2:**
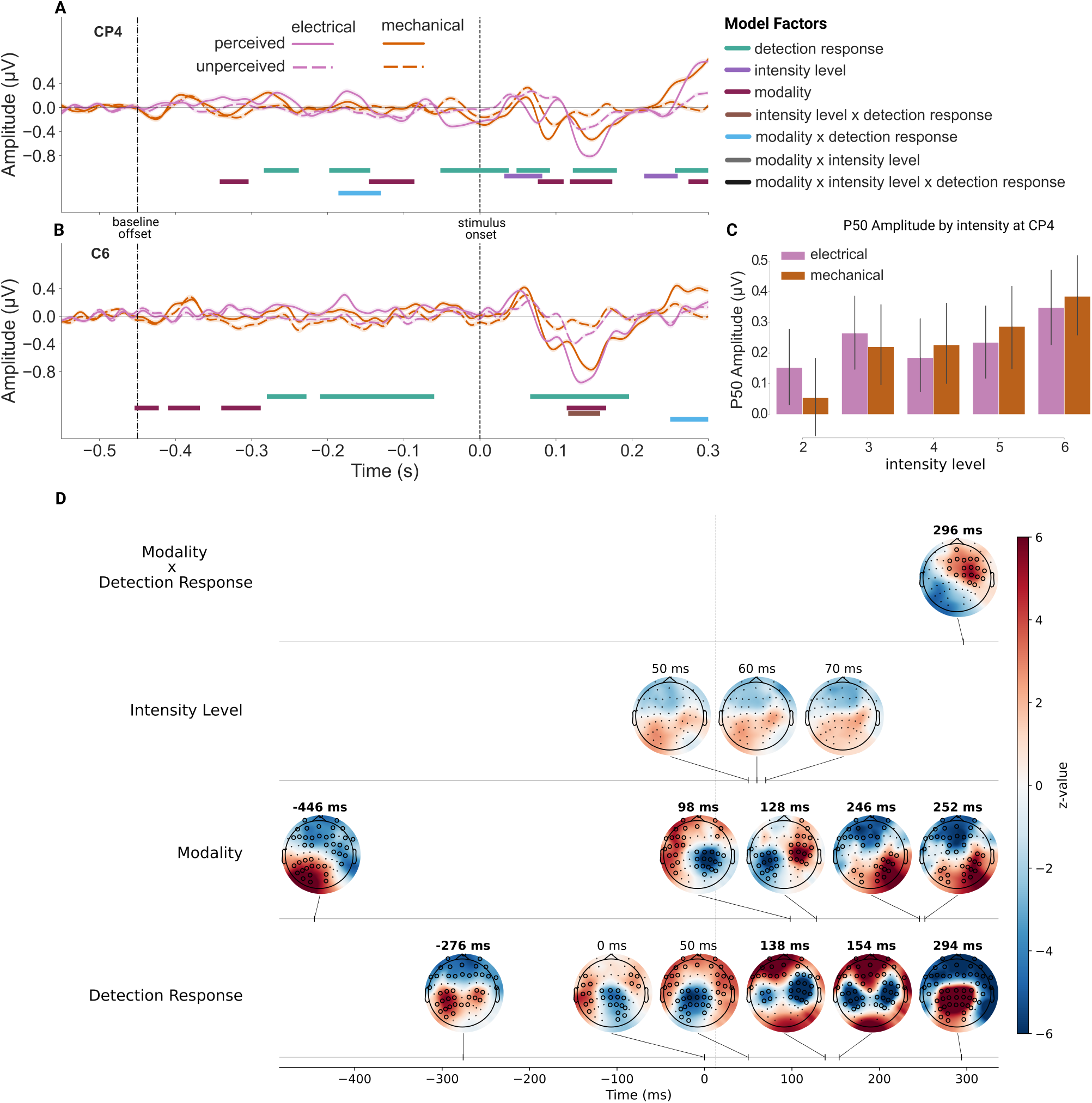
Analysis of the effect of detection response (perceived vs. unperceived), intensity level and modality on the event-related potential amplitude. Single channel analysis at A) CP4 and B) C6. In both A) and B) the time course of the measured ERP per modality and detection response is displayed. Colored bars indicate time points in significant clusters, according to the temporal cluster-based permutation analysis. Panel C) shows the effect of intensity on mean ERP amplitude in the time window 50-60ms post-stimulation at electrode CP4 (choice of time window and electrode based on [4–6]). D) Result of the full-electrode cluster-based permutation test displaying one row per model coefficient. Electrodes highlighted with a black circle belong to a cluster that indicates a significant difference for the specified factor. Topographies with time points highlighted in bold directly stem from cluster peak latencies, while non-highlighted ones are freely chosen latencies for demonstration. The interaction between response and intensity was not significant and is therefore not depicted here. The factor intensity level did not show any significant clusters in contrast to the single channel analysis, which is why their z-value topography is still depicted.

**Table 1:** Significant clusters of the single-channel temporal cluster-based permutation tests at CP4 and C6. At each channel the linear mixed model was fitted at every time point across both modalities over the interval *−*550 to 300 ms relative to stimulus onset. Two-sided cluster-mass permutation test along the time axis, 2000 permutations, cluster-forming threshold *z* = 1.96, corrected at *p* < .05. The mass is the sum of *z* values within the cluster. The peak is the sample of largest *|z|* inside the cluster, representing the effect size. Clusters are listed per factor in order of decreasing *|*mass*|*.

| Channel | Factor | Mass | Peak *z* | Peak *t*<br>(ms) | Time window<br>(ms) | *p* |
|---|---|---|---|---|---|---|
| CP4 | detection response | −167 | −7.58 | 146 | 122 to 180 | < .0005 |
|  |  | −166 | −4.98 | −32 | −52 to 38 | < .0005 |
|  |  | −137 | −9.29 | 74 | 48 to 92 | < .0005 |
|  |  | 128 | 8.50 | 300 | 256 to 300 | < .0005 |
|  |  | 107 | 4.83 | −168 | −198 to −144 | .001 |
|  |  | 80 | 4.11 | −266 | −284 to −238 | .006 |
|  | intensity level | 62 | 2.79 | 40 | 32 to 82 | .023 |
|  |  | −61 | −3.13 | 240 | 216 to 260 | .024 |
|  | intensity × response | *no significant cluster* |  |  |  |  |
|  | modality | 139 | 5.92 | 130 | 118 to 174 | < .0005 |
|  |  | −103 | −7.71 | 96 | 76 to 110 | .001 |
|  |  | 100 | 4.17 | −102 | −146 to −86 | .001 |
|  |  | −54 | −3.25 | −316 | −342 to −304 | .042 |
|  |  | −53 | −4.62 | 288 | 274 to 300 | .048 |
|  | modality × intensity | *no significant cluster* |  |  |  |  |
|  | modality × response | −68 | −2.56 | −154 | −186 to −130 | .009 |
|  | modality × intensity × response | *no significant cluster* |  |  |  |  |
| C6 | detection response | −468 | −11.86 | 140 | 66 to 196 | < .0005 |
|  |  | 254 | 5.07 | −180 | −210 to −60 | < .0005 |
|  |  | 86 | 3.93 | −246 | −280 to −228 | .001 |
|  | intensity level | *no significant cluster* |  |  |  |  |
|  | intensity × response | −56 | −2.84 | 136 | 116 to 158 | .019 |
|  | modality | 110 | 7.70 | 124 | 114 to 166 | < .0005 |
|  |  | −103 | −5.72 | −326 | −340 to −288 | < .0005 |
|  |  | 85 | 5.17 | −378 | −410 to −368 | .002 |
|  |  | −54 | −3.87 | −444 | −454 to −422 | .024 |
|  | modality × intensity | *no significant cluster* |  |  |  |  |
|  | modality × response | 86 | 4.81 | 290 | 250 to 300 | < .0005 |
|  | modality × intensity × response | *no significant cluster* |  |  |  |  |

### Broad band effects of stimulus detection and intensity pre- and post-stimulus

To reveal the topographies of the modality, detection response, and intensity effects we performed a spatiotemporal cluster-based permutation test (two-sided, 2000 permutations, corrected at *p* < .05) (see Fig. 2D and Table 2 for the complete statistical results). Significant clusters emerged for the factors detection response and modality, and for their interaction; stimulus intensity did not yield any cluster. All significant clusters are listed in Table 2.

**Table 2:** Significant clusters of the full spatio-temporal cluster-based permutation test. The linear mixed model was fitted across both modalities (40 participants, 74 sessions: 35 mechanical, 39 electrical) over the interval *−*550 to 300 ms relative to stimulus onset on all 62 channels. Two-sided cluster-mass permutation test, 2000 permutations, cluster-forming threshold *z* = 1.96, corrected at *p* < .05. Mass is the sum of *z* values within the cluster; its sign gives the cluster’s polarity. The peak is the sample of largest *|z|* inside the cluster; time window and number of channels give its temporal and spatial extent. Clusters are listed per factor in order of decreasing *|*mass*|*.

| Factor | Mass | Peak *z* | Peak ch. | Peak *t*<br>(ms) | Time window<br>(ms) | Ch.<br>(*n*) | *p* |
|---|---|---|---|---|---|---|---|
| detection response | −14948 | −12.87 | C4 | 138 | −198 to 300 | 62 | < .0005 |
|  | 13516 | 9.26 | FPz | 154 | −384 to 202 | 60 | < .0005 |
|  | 2415 | 11.34 | CP2 | 294 | 240 to 300 | 27 | < .0005 |
|  | −2148 | −5.26 | FPz | −276 | −378 to −236 | 24 | < .0005 |
| intensity level | *no significant cluster* |  |  |  |  |  |  |
| intensity × response | *no significant cluster* |  |  |  |  |  |  |
| modality | −4566 | −5.73 | C6 | −326 | −474 to −212 | 61 | < .0005 |
|  | −2888 | −7.67 | CP2 | 98 | 70 to 196 | 53 | < .0005 |
|  | 2759 | 8.03 | C4 | 128 | 66 to 178 | 57 | < .0005 |
|  | 2635 | 5.93 | O2 | −46 | −56 to 66 | 56 | < .0005 |
|  | −2360 | −5.84 | FT10 | −4 | −86 to 66 | 58 | < .0005 |
|  | 1962 | 5.19 | C4 | −98 | −218 to −80 | 38 | < .0005 |
|  | −1869 | −6.14 | F2 | 252 | 224 to 300 | 33 | < .0005 |
|  | 1636 | 7.00 | P8 | 246 | 194 to 300 | 24 | .002 |
|  | −1469 | −5.23 | FT10 | −120 | −170 to −84 | 31 | .004 |
|  | 1215 | 5.37 | O2 | −236 | −286 to −220 | 23 | .012 |
|  | 1182 | 5.84 | AF8 | −414 | −428 to −356 | 32 | .015 |
|  | 919 | 6.95 | O1 | −446 | −470 to −426 | 20 | .049 |
| modality × intensity | *no significant cluster* |  |  |  |  |  |  |
| modality × response | 1173 | 5.44 | C4 | 296 | 192 to 300 | 25 | .016 |
| modality × intensity × response | *no significant cluster* |  |  |  |  |  |  |

The factor detection response (perceived vs. unperceived) revealed four clusters in total, spanning time points before and after the stimulus onset. Already in the cue-to-stimulus interval a negative cluster over 24 frontal electrodes was present (peak *z* = *−*5.26 at FPz at -276 ms, *p* < .0005). After stimulus onset the effect was far stronger and at some times spanned all sensors. A negative cluster covered all 62 electrodes with a peak at C4 at 138 ms (*z* = *−*12.87, *p* < .0005) together with a positive cluster over 60 electrodes with a peak at FPz at 154 ms (*z* = 9.26, *p* < .0005). Since both peak at nearly the same latency but with opposite polarity and complementary topographies, they might represent the same neural process rather than two independent effects near their peak latencies. A fourth, later positive cluster comprised 27 centro-parietal and occipital electrodes between 240 and 300 ms (peak *z* = 11.34 at CP2 at 294 ms, *p* < .0005).

The factor modality (mechanical vs. electrical) produced twelve clusters that together covered the entire tested interval, from -474 ms up to the end of the epoch. The two strongest peaks were again of opposite polarity and centred on the early somatosensory response: a negative cluster over 53 electrodes with a peak at CP2 at 98 ms (*z* = *−*7.67, *p* < .0005) and a positive cluster over 57 electrodes with a peak at C4 at 128 ms (*z* = 8.03, *p* < .0005). Modality also differed within the pre-stimulus interval, with a widespread negative cluster from -474 to -212 ms (the cluster with the largest mass; 61 channels, peak *z* = *−*5.73 at C6 at -326 ms, *p* < .0005) and a positive cluster from -218 to -80 ms (38 channels, peak *z* = 5.19 at C4 at -98 ms, *p* < .0005). Further clusters of opposite polarity surrounded stimulus onset, from -56 to 66 ms (positive, 56 channels, peak *z* = 5.93 at O2 at -46 ms) and from -86 to 66 ms (negative, 58 channels, peak *z* = *−*5.84 at FT10 at -4 ms), and late clusters at the end of the epoch from 194 to 300 ms (positive, 24 channels, peak *z* = 7.00 at P8 at 246 ms, *p* = .002) and from 224 to 300 ms (negative, 33 channels, peak *z* = *−*6.14 at F2 at 252 ms, *p* < .0005). The remaining pre-stimulus clusters were smaller (Table 2).

The interaction between modality and detection response reached significance only late and weakly, in one positive cluster over 25 right fronto-central electrodes from 192 to 300 ms, which extended to the end of the epoch (peak *z* = 5.44 at C4 at 296 ms, *p* = .016). The detection effect is therefore largely comparable across the two modalities within the tested window, and diverges only in the late post-stimulus interval. Stimulus intensity yielded no significant cluster, and neither did its interaction with detection response or its interaction with modality.

### Neural origin of significant cluster topographies

We performed source reconstruction of the *z* value topographies for visual interpretation of significant clusters (see Methods). We projected the group-level *z* topography onto the cortical surface, averaged over the 57 sessions with an available T1 scan (28 mechanical, 29 electrical), and displayed it as absolute source amplitude (Fig. 3). For detection response, the estimate in the cue interval, at the peak of the negative frontal sensor cluster (-276 ms; 24 electrodes, -378 to -236 ms), was mainly left-hemispheric (74% left, 26% right) and lay in the left postcentral (28%) and precentral gyrus (27%) and the right postcentral gyrus (16%). The strongest source vertices of this frontal sensor cluster were therefore sensorimotor and not frontal. The estimate was focal and clearly lateralised at the peak of the largest and strongest post-stimulus cluster (138 ms): all of the strongest 1% of vertices lay in the right hemisphere, contralateral to the stimulated hand, and were concentrated in the right precentral (39%) and postcentral gyrus (37%), extending into the right supramarginal gyrus (11%). At the peak of the positive cluster (154 ms), the opposite pole of the same dipolar sensor field, 59% of the vertices were left-hemispheric (left postcentral 27%, left supramarginal 9%) and 41% right-hemispheric (right postcentral 9%, right superior temporal 9%). At 294 ms the focus returned to the right hemisphere (60% right, right precentral 40%, right postcentral 20%; left postcentral 22%, left precentral 17%). Immediately around the stimulus onset we observed a particular centred topography in the significant clusters (though there is no cluster peak at these latencies). At stimulus onset we found a bilateral pattern centred on the pre- and postcentral gyri (57% right and 43% left; right precentral 33%, left precentral 22%, right postcentral 20%). At 50 ms the focus was on the left hemisphere (74% left-hemispheric; left precentral 42%, left postcentral 29%; 26% right, right postcentral 13%). Both place the strongest vertices in the pre- and postcentral gyri, but the right-hemispheric share is larger at 0 ms (57%) than at 50 ms (26%).

**Figure 3:**
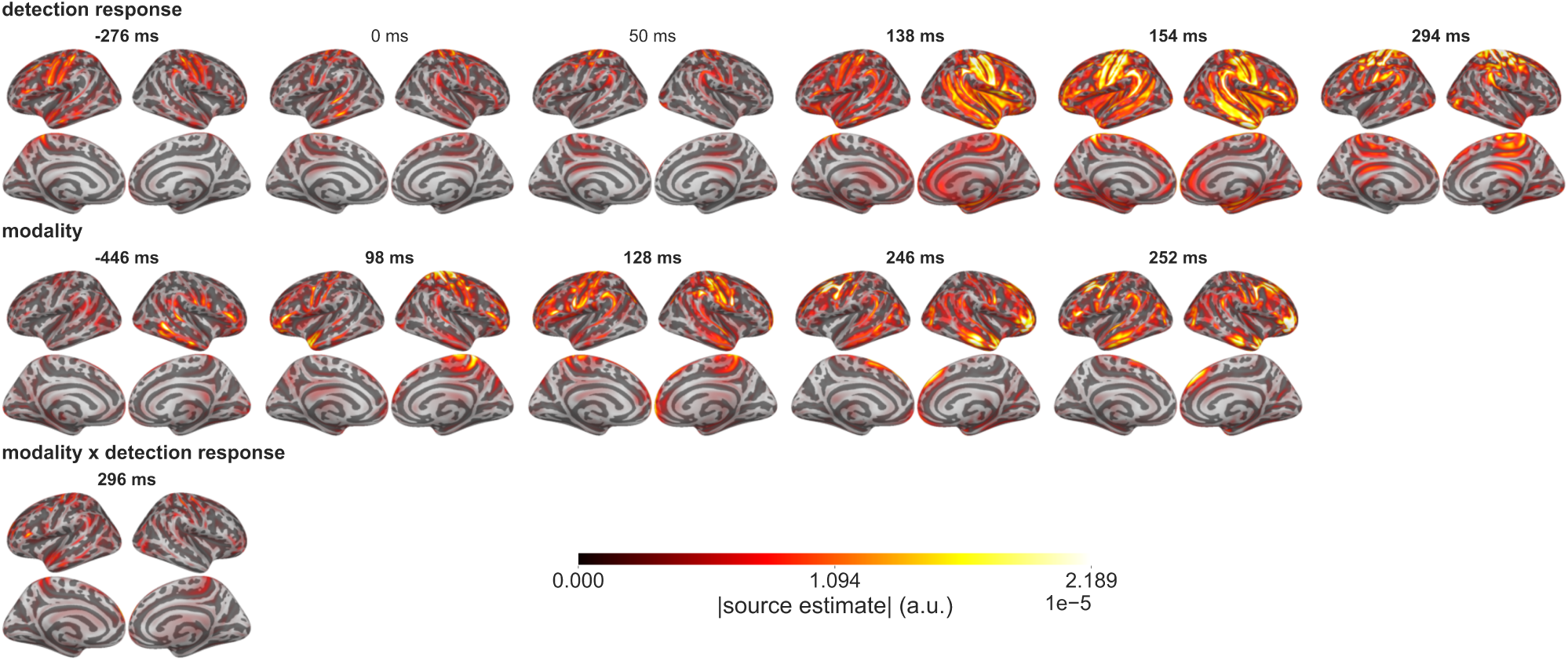
Source reconstruction of the group-level *z* value topographies of the three coefficients carrying the strongest significant sensor-level clusters, detection response (top) and modality (middle), modality x detection response (bottom). Each panel shows both hemispheres in lateral (upper row) and medial (lower row) view, with the left hemisphere on the left; latencies are given above the columns and include the peaks of the clusters with the highest z-values of each factor (98 ms for modality, 138 ms for detection response) as plotted in sensor space in Fig. 2D. Colour codes the absolute source amplitude on a scale shared by both factors and all latencies. Time points highlighted in bold directly stem from cluster peak latencies, while non-highlighted ones are freely chosen latencies for demonstration. Note that the maps are descriptive and were not tested statistically in source space.

For modality we reconstructed the sources for the 5 clusters that showed the peaks with the highest z-values. Before the stimulus (-446 ms) the strongest vertices were mostly right-hemispheric (86% right, 14% left) and lay in the right temporal cortex (middle temporal 20%, inferior temporal 10%) and right inferior frontal cortex (pars triangularis 10%), with the left hemisphere contributing the left lateral occipital gyrus (10%). At the early evoked response (98 ms) the modality effect engaged the same contralateral sensorimotor region as in the detection effect (86% right-hemispheric, right postcentral 30%, right precentral 29%, right superior parietal 12%), with a smaller left-hemispheric part (14%) in the left superior temporal gyrus (9%). The effect was more widely distributed than the detection effect and shifted its focus over time. At 128 ms it was still right lateralised but weaker, with 60% right and 40% left hemispheric activation (right postcentral 23%, right rostral middle frontal 11%, left postcentral 10%, left precentral 9%, left superior frontal 8%). After 200 ms the focus moved away from sensorimotor cortex towards frontal cortex in both hemispheres (246 ms: 55% right and 45% left; left caudal middle frontal 21%, left precentral 17%, right middle temporal 16%, right superior frontal 11%; 252 ms: 75% right and 25% left; left caudal middle frontal 13%, right superior frontal 12%, right rostral middle frontal 11%, right pars orbitalis 11%, right caudal middle frontal 11%).

In summary, the effect of stimulus detection showed a focal contralateral sensorimotor generator around the post-stimulus peak at 138 ms, with an additional left-hemispheric focus shortly afterwards, whereas the modality effect reflects a broader network that involves right temporal and inferior frontal cortex before the stimulus and frontal cortex late in the epoch. The modality *×* detection response interaction (296 ms) was left-lateralised (70% left, 30% right) and spread over several regions (left pre-central 14%, right postcentral 12%, left rostral middle frontal 11%, right precentral 9%, left postcentral 9%).

### Alpha power in mechanical and electrical stimulation affects stimulus detection

Pre-stimulus alpha envelope was extracted from the surface-Laplacian transformed epochs over the pre-stimulus window (-400 to -100 ms) and sorted into ten equiprobable bins per subject (36 subjects in the electrical and 32 in the mechanical modality after the SNR-based exclusion). The quadratic coefficient of a second-order polynomial fitted to each subject’s bin-wise detection rate was negative at every electrode. Testing this coefficient one-sided against zero (H_1_: *a <* 0) and FDR-correcting across the three channels within each modality, the effect was significant in the electrical modality at C4 (*a* = *−*1.26 *×* 10*^−^*^3^, *p*_FDR_ = .009), but not at O2 (*a* = *−*6.6 *×* 10*^−^*^4^, *p*_FDR_ = .113) or CP4 (*a* = *−*4.7*×*10*^−^*^4^, *p*_FDR_ = .158). In the mechanical modality it was significant at O2 (*a* = *−*1.26*×*10*^−^*^3^, *p*_FDR_ = .021), but did not survive the correction at the somatosensory electrodes (C4: *a* = *−*7.9*×*10*^−^*^4^, *p* = .050, *p*_FDR_ = .057; CP4: *a* = *−*7.1 *×* 10*^−^*^4^, *p*_FDR_ = .057). The inverted-U relationship is thus significant in the electrical modality only at the somatosensory electrode C4, and in the mechanical modality only at the occipital control site.

We compared the mean pre-stimulus alpha envelope directly between perceived and unperceived trials for each modality separately (Fig. 4B). Alpha envelope was lower before detected than before undetected trials at both contralateral somatosensory electrodes in the electrical modality (CP4: *t*(35) = *−*3.28, *p*_FDR_ = .004; C4: *t*(35) = *−*3.60, *p*_FDR_ = .003). In the mechanical modality the difference went in the same direction but was not significant (C4: *t*(31) = *−*2.23, *p*_FDR_ = .100; CP4: *t*(31) = *−*1.51, *p*_FDR_ = .21). At the occipital electrode O2 the mean envelope did not differ between perceived and unperceived trials in either modality (electrical: *t*(35) = *−*1.35, *p*_FDR_ = .19; mechanical: *t*(31) = *−*0.21, *p*_FDR_ = .83).

**Figure 4:**
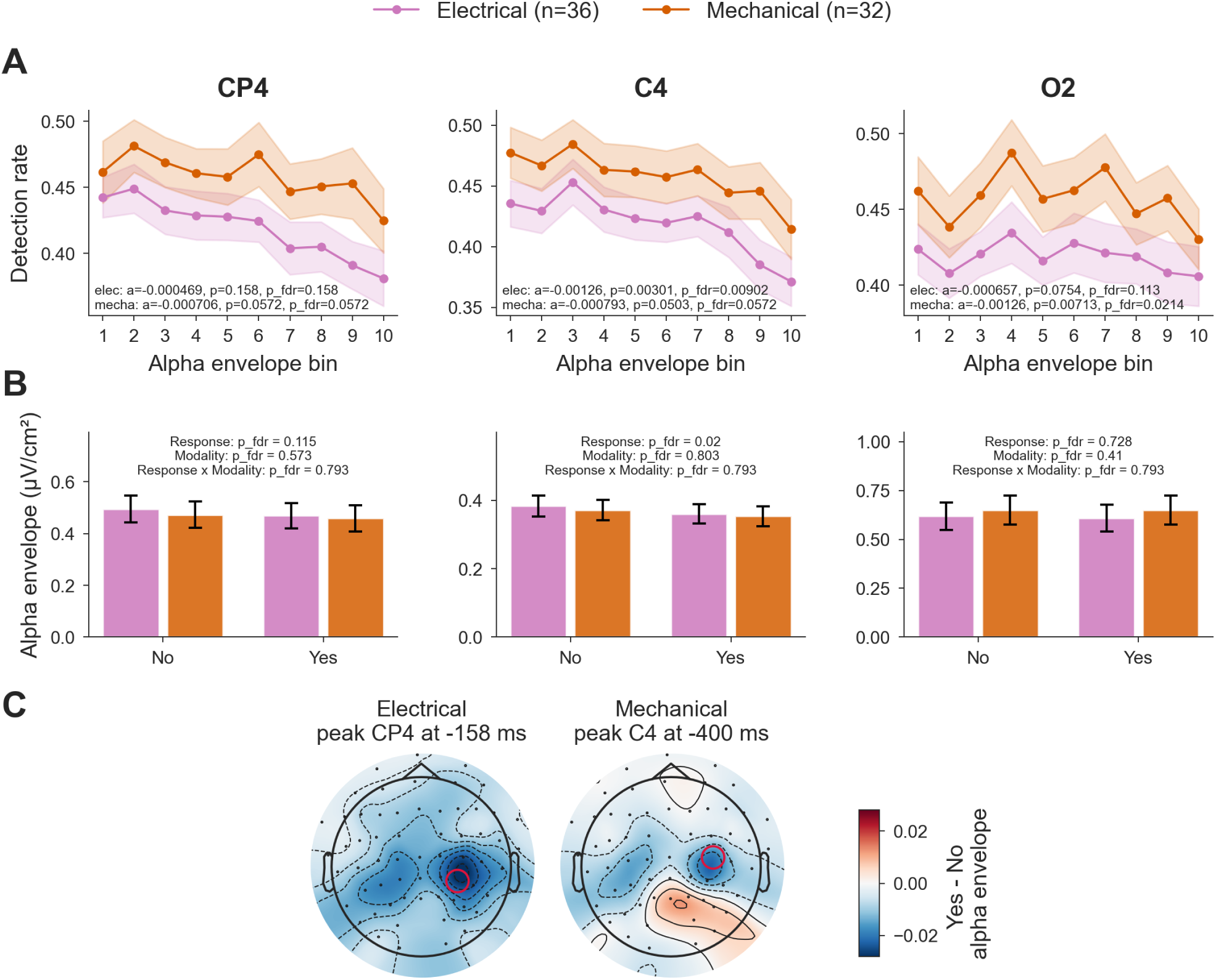
Effects of pre-stimulus alpha envelope on detection rate in both modalities. A) Detection rate by alpha bin (higher bin number means higher envelope amplitude) for three channels CP4, C4 and O2. Statistics show the quadratic effect of alpha envelope on the detection rate. Alpha bins are computed within-subject. The shaded bands indicate the standard error of the mean between subjects. B) Alpha envelope per detection response. Statistics in A) and B) are FDR-corrected (Benjamini-Hochberg) across the three tested channels within each modality. The text in B) gives the FDR-corrected p-values of the response *×* modality linear mixed model (main effects and interaction). C) Topography showing the strongest (red highlighted electrode) difference in alpha envelope comparing perceived and unperceived trials.

To compare the modalities directly, we fitted a linear mixed model to the subject-level envelope. At C4 the envelope was lower for detected trials across both modalities (*β* = *−*0.021, SE = 0.008, *z* = *−*2.71, *p*_FDR_ = .020). At CP4 this effect did not survive correction (*β* = *−*0.020, SE = 0.011, *z* = *−*1.77, *p*_FDR_ = .115), and at O2 there was none (*p*_FDR_ = .73). Neither the main effect of modality nor the response *×* modality interaction was significant at any electrode (all *p*_FDR_ *≥ .*41), so the weaker mechanical effect did not differ significantly from the electrical one.

The topography of the Yes-No alpha envelope difference was negative over most of the scalp in both modalities, i.e. reduced pre-stimulus alpha amplitude preceded detected trials (Fig. 4C). For electrical stimulation the difference was maximal at CP4 at -158 ms, and for mechanical stimulation the strongest deflection fell at C4 at -400 ms, the beginning of the analysis window. Given the weak and non-significant mechanical envelope effect at the somatosensory electrodes, this peak at C4 should be interpreted with caution.

### Phase locking in the alpha band as preparation for stimulus detection

We examined the dITPC before and after the visual cue, after regressing the absolute broadband ERP out of the dITPC time course, so that the reported effects are not explained by the evoked response. In electrical stimulation, dITPC at electrode CP4 was significantly higher for perceived compared to unperceived trials in the post-cue window (*t*(33) = 3.85, *p*_FDR_ = .001, *d* = 0.66), indicating that stronger phase locking to the stimulus cue is associated with subsequent stimulus detection (Fig. 5A).

**Figure 5:**
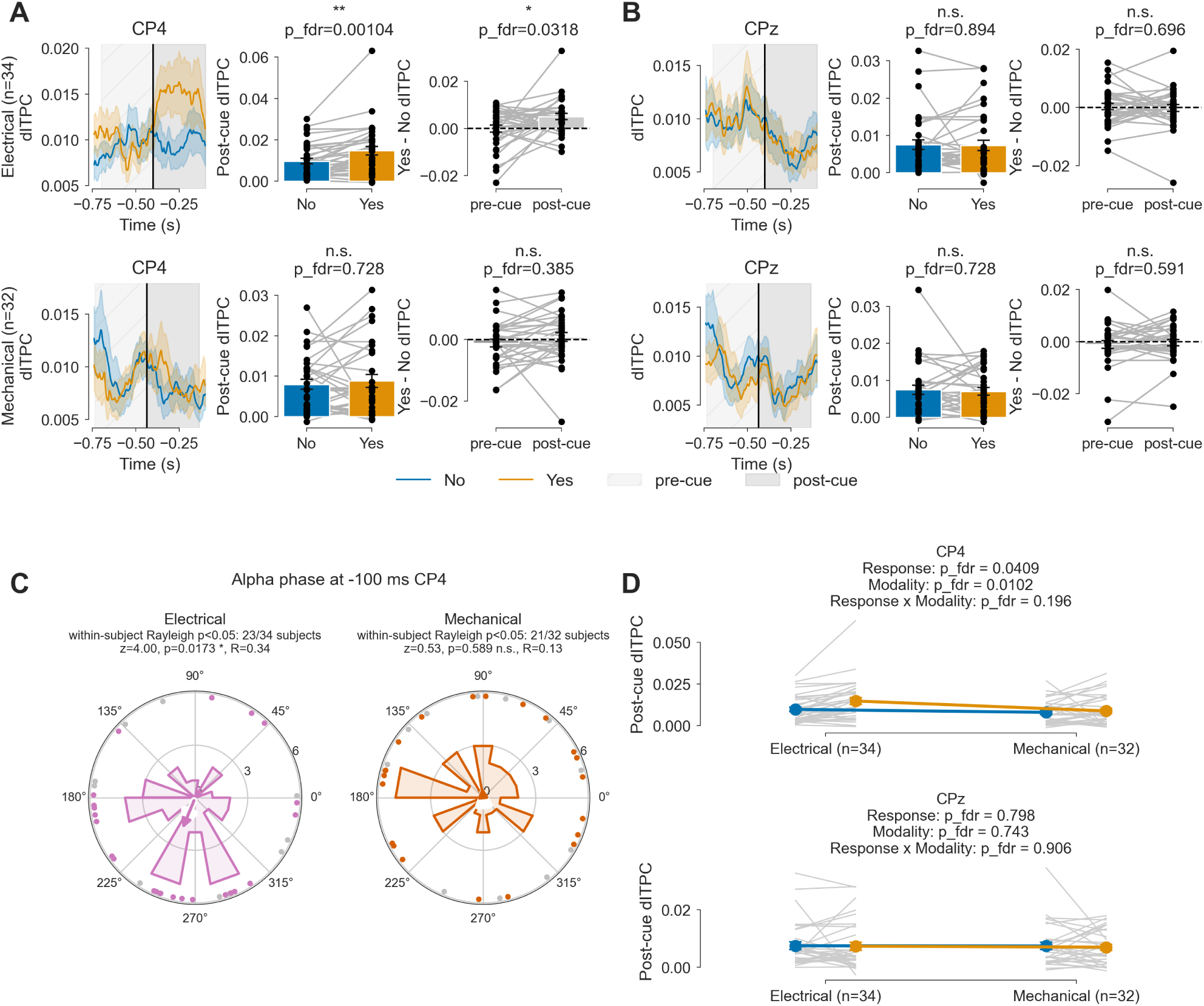
Effect of inter-trial phase coherence (dITPC) on detection outcome, after regressing the absolute broadband ERP out of the dITPC time course. A) and B) show the results for channel CP4 and CPz, each with the dITPC time course (perceived vs unperceived), the mean post-cue dITPC and the Yes-No dITPC difference before vs after cue onset, separately for the electrical (top) and mechanical (bottom) modality. Statistics are FDR-corrected (Benjamini-Hochberg) across the two tested channels within each modality. C) Phase of the alpha oscillation at 100 ms pre-stimulus at CP4, one circular mean per subject, pooled perceived and unperceived trials. Rayleigh test conducted per modality, uncorrected. Subjects whose single-trial phases at this time point were not consistent across trials (Rayleigh test within subject, *p ≥ .*05, uncorrected) are shown as grey dots; the number of subjects with a significant test is given under each modality name. D) Post-cue dITPC per subject (grey lines connect the perceived and unperceived value of one subject), cell mean *±* SEM (coloured markers) by detection response and modality at CP4 (top) and CPz (bottom). The statistics below each channel name give the FDR-corrected p-values of the main effects of response and modality and of the response *×* modality interaction (linear mixed model, FDR-corrected across the two channels per effect).

This effect was not significant at the central electrode CPz (*t*(33) = *−*0.13, *p*_FDR_ = .89, Fig. 5B), nor in the mechanical modality at either electrode (CP4: *t*(31) = 0.59, *p*_FDR_ = .73; CPz: *t*(31) = *−*0.35, *p*_FDR_ = .73). The Yes-No dITPC difference at CP4 in the electrical modality was larger after cue onset than in the pre-cue window (*t*(33) = 2.54, *p* = .016, *p*_FDR_ = .032, *d* = 0.44). The pre- to post-cue change was absent at CPz (*t*(33) = *−*0.39, *p*_FDR_ = .70) and not significant in the mechanical modality (CP4: *t*(31) = 1.33, *p*_FDR_ = .38; CPz: *t*(31) = 0.54, *p*_FDR_ = .59). The effect is therefore restricted to the cue-to-stimulus interval in electrical stimulation at CP4.

To compare the post-cue dITPC effect directly between the modalities, we fitted a linear mixed model to the subject-level post-cue dITPC (Fig. 5D; *n* = 38 subjects, of which 34 electrical and 32 mechanical subject-sessions; 28 subjects contributed both sessions). At CP4, dITPC was higherfor perceived than for unperceived trials (response: *β* = 0.0030, SE = 0.0013, *z* = 2.32, *p* = .020, *p*_FDR_ = .041) and higher in the electrical than in the mechanical modality (modality: *β* = *−*0.0037, SE = 0.0013, *z* = *−*2.80, *p* = .005, *p*_FDR_ = .010). The response *×* modality interaction was not significant (interaction: *β* = *−*0.0042, SE = 0.0025, *z* = *−*1.66, *p* = .098, *p*_FDR_ = .20). At CPz, neither main effect nor the interaction reached significance (response: *β* = *−*0.0003, SE = 0.0011, *z* = *−*0.26, *p*_FDR_ = .80; modality: *β* = *−*0.0004, SE = 0.0012, *z* = *−*0.33, *p*_FDR_ = .74; interaction: *β* = *−*0.0003, SE = 0.0023, *z* = *−*0.12, *p*_FDR_ = .91).

To examine whether there is a specific alpha phase right before the stimulus, independent of the detection outcome, we computed the alpha phase 100 ms pre-stimulus (Fig. 5C). In electrical stimulation, the alpha phase concentrated around a preferred angle (Rayleigh test, mean phase *≈ −*112*^◦^*, *R* = 0.34, *z* = 4.00, *p* = .017, *n* = 34, uncorrected). This was not the case for mechanical stimulation, where the alpha phase seemed rather randomly distributed (Rayleigh test, *R* = 0.13, *z* = 0.53, *p* = .59, *n* = 32). As a reliability check of the subject-level phases, we additionally tested for each session whether the single-trial phases at -100 ms were consistent across trials (Rayleigh test within session, *p* < .05, uncorrected). This was the case in 23 of 34 electrical and 21 of 32 mechanical sessions.

To locate the dITPC effect we applied a two-sided spatio-temporal cluster-based permutation test to the per-subject Yes-No difference maps of the source reconstructed alpha band activity, separately for each modality (2000 permutations, cluster-forming threshold *t* = 2.06, -750 to -100 ms; 26 subjects in each modality). No cluster reached significance in either modality (smallest cluster-level *p* = .54 for electrical and *p* = .52 for mechanical stimulation), so the sensor-level dITPC effect at CP4 could not be localised in source space.

### Interactions between alpha effects, P50, and preparation negativity

We investigated if the alpha phase effect related to the early P50 post-stimulus evoked response. The P50 at CP4 increased with the pre-stimulus alpha amplitude bin in the mechanical, but not in the electrical modality (Fig. 6A). Fitting one line per session across the eight bins and testing the slopes against zero gave a positive effect for mechanical stimulation (slope = 0.041 *µ*V per bin, *t*(31) = 3.15, *p* = .004), indicating that higher pre-stimulus alpha amplitude preceded a larger early evoked response. The correlation between alpha power bin and P50 amplitude was not significant in the electrical condition (slope = 0.030 *µ*V per bin, *t*(33) = 1.92, *p* = .064). Split by detection response (uncorrected), the mechanical slope was significant for unperceived trials (slope = 0.056 *µ*V per bin, *t*(31) = 2.37, *p* = .024) but not for perceived trials (slope = *−*0.005 *µ*V per bin, *t*(31) = *−*0.13, *p* = .90); in the electrical modality neither response reached significance (perceived: slope = 0.034 *µ*V per bin, *t*(33) = 1.58, *p* = .124; unperceived: slope = 0.021 *µ*V per bin, *t*(33) = 1.00, *p* = .33).

**Figure 6:**
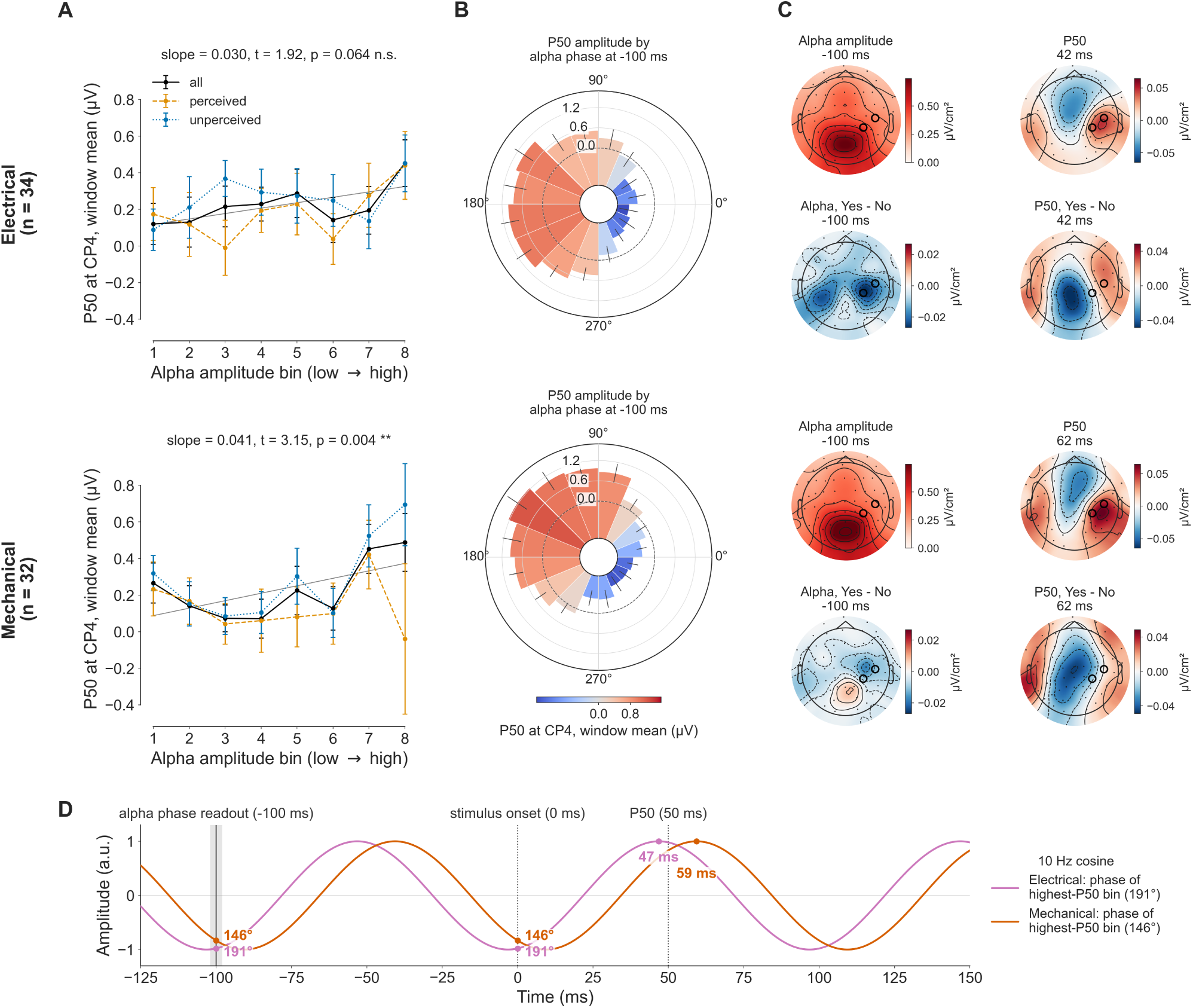
Effect of pre-stimulus alpha amplitude and phase on the P50. Rows show the electrical (top) and the mechanical (bottom) modality. A) P50 amplitude at CP4 across eight equiprobable pre-stimulus alpha amplitude bins (higher bin number means higher envelope amplitude), pooled over responses (black) and split into perceived (orange) and unperceived (blue) trials; error bars give the standard error across sessions, the grey line the linear trend, and the title the session-level slope test. B) P50 amplitude at CP4 by pre-stimulus alpha phase at -100 ms, in 16 bins of equal angular width. Bar length and bar colour both encode the bin mean P50; the dashed circle marks 0 *µ*V. Whiskers give the standard error across sessions. C) Topography comparison of pre-stimulus alpha activity and P50. The alpha topography represents the absolute Hilbert envelope at 100 ms pre-stimulus. The P50 topography is obtained at a fixed latency per modality (42 ms electrical, 62 ms mechanical). Prior to computing the difference topography we normalized the maps across electrodes by computing the root mean square of the centered map. D) Illustration of a 10 Hz alpha cycle (cosine convention: 0° = peak, 180° = trough) with the phase at -100 ms set to the strongest phase bin of electrical and mechanical trials at CP4.

Sorting the trials by the alpha phase at -100 ms instead of by amplitude produced a graded pattern across the circle in both modalities (Fig. 6B). The bin-wise P50 was largest for phase bins in the second and third quadrants, peaking in the 180°-202.5° bin for the electrical modality (1.02 *µ*V) and in the 135°-157.5° bin for the mechanical modality (1.23 *µ*V).

To test whether the P50 reflects a topography distinct from the pre-stimulus alpha oscillation, we compared the pre-stimulus alpha and P50 topographies using TANOVA (Fig. 6C). The observed global dissimilarity was significantly larger than expected under 5000 sign-flip permutations for both electrical (DISS = 0.94, null 95th percentile = 0.40, *p* = .0002) and mechanical stimulation (DISS = 0.95, null 95th percentile = 0.39, *p* = .0002), confirming that the P50 and pre-stimulus alpha topographies differ significantly in both modalities. The perceived versus unperceived difference topographies of alpha and P50 also differed in shape in both modalities (electrical: DISS = 0.41, null 95th percentile = 0.31, *p* = .0008; mechanical: DISS = 0.46, null 95th percentile = 0.34, *p* = .0002; Fig. 6C, bottom row of each block).

Finally, we tested whether pre-stimulus phase locking relates to the amplitude of the negative preparation component at CPz (Fig. 7). Since the component is measured as a negative peak, a negative correlation means that stronger phase locking accompanies a larger preparation negativity.

**Figure 7:**
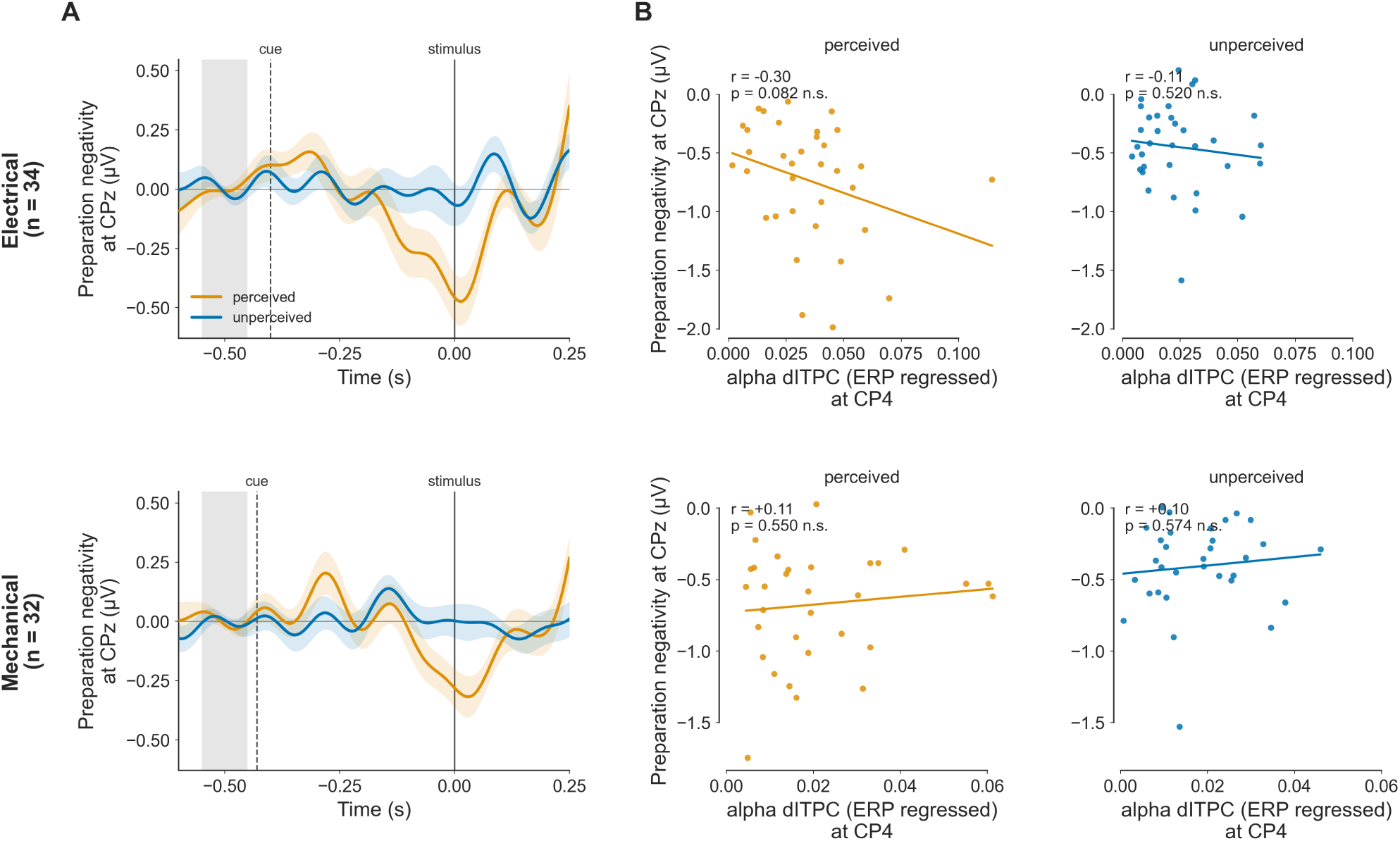
A) The time course of the preparatory negativity signal by detection response and modality. B) Correlation between alpha dITPC and the amplitude of the preparatory negativity signal by detection response.

None of the four correlations was significant after FDR correction (all *p*_FDR_ *≥ .*33). The strongest relationship was a negative correlation for perceived trials in the electrical modality (*r* = *−.*30, *p* = .082, *p*_FDR_ = .33, *n* = 34). The remaining three combinations were flat (electrical unperceived: *r* = *−.*11, *p* = .52, *n* = 34; mechanical perceived: *r* = .11, *p* = .55, *n* = 32; mechanical unperceived: *r* = .10, *p* = .57, *n* = 32).

## Discussion

In this study, we investigated pre- and post-stimulus processes during mechanical and electrical so-matosensory perception. We found potential correlates of conscious perception in a time window that included the N80 and the N140 at the electrode C6 in our single channel analysis. Most studies reported the N140 as a neural correlate of consciousness [5, 6, 43], while some studies reported earlier differences between perceived and un-perceived stimulation [4, 10, 19]. The spatio-temporal cluster-based permutation analysis revealed the earliest post-stimulus cluster peak at 138 ms at electrode C4, which matches the N140 latency [2, 6]. In the spatio-temporal analysis we observed a strong fronto-central negative preparation effect starting already before stimulation onset. It was more negative for perceived compared to unperceived trials, which was also visible in the single-channel analysis of the electrode CP4 showing a lower amplitude from 52 ms before stimulation until 38 ms post-stimulus. Importantly, there was no interaction effect of modality and stimulus detection at the latencies of the P50, N80 or N140, as we would have expected based on the findings by Förster et al. [4], who reported a fully absent P50 component for unperceived trials in the mechanical condition. Interaction clusters appeared only later in the epoch: in the spatio-temporal analysis in one right fronto-central positive cluster from 192 ms (peak at C4 at 296 ms) that lasted until the end of the epoch. In the single channel analysis of CP4 we observed an interaction even before the stimulus (-186 to -130 ms), suggesting different preparation mechanisms in the broad band activity. Instead, we found an effect of stimulus intensity at 32 to 82 ms, which is characteristic of the P50 component and in line with previous literature showing that the P50 reflects processing of physical characteristics of the stimulus [5, 6]. One might argue that because the ERP trace observed in channel CP4 is contaminated by strong oscillatory signals already before stimulus onset, early post-stimulus responses are just a continuation of those oscillations. When examining the topographies of the P50 component at the respective time points per modality, we observed a clear lateral dipole (Fig. 6C) with a profile that is highly similar to what was previously reported as a P50 topography [5, 6]. The electrode CP4 was closer to the hotspot of pre-stimulus occipital alpha and therefore its trace showed more contamination than the electrode C6, where we observed a clear P50 peak for both modalities. Besides that, we observed a delayed mechanical P50 peak (average peak of 62 ms) compared to the electrical P50 (average peak of 42 ms) and similarly for later component peaks, which is in line with Alouit et al. [44]. They directly compared supra threshold electrical and mechanical stimulation and observed increasingly higher delays in ERP component peaks for the mechanical modality. One possible cause for the absence of the P50 in the study by Förster et al. [4] could be the lack of preparation towards the stimulus. Expectancy might induce effects that could overlap with a weaker effect early post-stimulus. Another reason for the diverging findings might be the difference in performing the threshold assessment. Jones et al. [10] reported a significantly larger component peak at 75 ms for a static threshold, but not when using a dynamically adapting stimulus threshold. Gusso et al. [11] applied vibrotactile stimulation and used a dynamical assessment method by continuously maintaining the threshold at 50% detection probability. They found no statistically significant ERP components in the 50–70 ms range. We performed threshold assessment before each experimental block (8 times in total), while Förster et al. [4] performed only one assessment before the experiment.

We did find an effect of the main factor modality in the N140 window in C6 as well as significant spatio-temporal clusters, which spanned the entire epoch and nearly all electrodes. In the full channel analysis, the positive modality cluster had its maximum at 128 ms post-stimulus at the electrode C4 (*z* = 8.03). In source space, the right hemisphere dominated, but only marginally with 60% right and 40% left activation of the strongest vertices (right postcentral gyrus 23%, right rostral middle frontal gyrus 11%, left postcentral gyrus 10%, left precentral gyrus 9%). Earlier in time, at 98 ms, the pre- and postcentral gyrus corresponded more clearly to the modality effect. Together this suggests different pathways for the two modalities during that time window, in line with a previous study showing connectivity pattern differences between mechanical and electrical stimulation at mid- to late-latency stages [44]. Due to the issue that the preparation cue was presented 36 ms earlier in the mechanical condition (see Methods section), the early pre-stimulus clusters should be interpreted with caution. Furthermore, we observed an interaction between intensity and detection at C6 during the N140. For perceived trials, the effect of intensity on the amplitude was negative, meaning stronger intensities evoked a more negative N140 component. In contrast, when the stimulus was not perceived, the amplitude increased with stimulation intensity. This indicates that the N140 still transfers some information on stimulus characteristics, and even when the stage of conscious processing was not achieved, this information does not seem to be lost.

The source reconstruction of the statistical maps revealed post-stimulus activation exclusively in the right hemisphere, in the pre- and post-central gyrus at the peak of the detection effect at 138 ms. This speaks for contributions of S1 towards a conscious percept, which is in line with [10, 43, 45] but does not let us conclude if S1 is sufficient for perception. The left hemisphere contributed more strongly shortly afterwards, at 154 ms (59% left-hemispheric, left postcentral gyrus 27%). The precentral gyrus includes premotor cortex, whose involvement in somatosensory perception was previously reported by de Lafuente and Romo [46], who conducted a vibrotactile near-threshold experiment in macaque monkeys. They discovered that premotor activity increasingly correlated with stimulus perception and ruled out a simple motor preparation effect. It is still an ongoing debate which brain region is sufficient for conscious perception, which is impossible to answer with a purely correlational experimental design. We found that brain activity in somatosensory electrodes differs significantly between perceived and unperceived trials already before stimulation onset, starting at 284 ms pre-stimulus at electrode CP4 and at 280 ms in electrode C6 (see Fig. 2). The spatio-temporal analysis over all channels revealed a frontal negative cluster (-378 to -236 ms) and a broad positive cluster starting at -384 ms. The source estimate at the peak of the frontal cluster (-276 ms) was mainly left-hemispheric (74%) and lay in the left postcentral and precentral gyrus, so the frontal sensor cluster does not point to a frontal generator.

The ipsilateral hemisphere therefore seems to play a role in the preparation for stimulus perception.

### Alpha phase coherence and amplitude affect stimulus detection

We found that phase locking in the alpha band before stimulus onset might aid somatosensory perception.

In electrical stimulation, the alpha phase concentrated at a specific angle within the pre-stimulus time window, with a significant Rayleigh test at -100 ms (*p* = .017, Fig. 5C). When separately looking at perceived and unperceived trials, an increase in pre-stimulus alpha-band inter-trial phase coherence (dITPC) before stimulus onset could be observed, indicating that better tuning results in higher chances of perceiving the stimulus. This effect was observed at electrode CP4, located closest to the contralateral primary somatosensory cortex, whereas no comparable increase in pre-stimulus dITPC was found at the central electrode CPz. The difference in phase coherence between perceived and unperceived trials was significantly larger after cue onset than before it, suggesting that the cue plays a role in a phase-alignment process that facilitated subsequent stimulus detection.

The mechanical modality exhibited no significant pre-stimulus alpha phase locking and did not point to a specific phase at 100 ms pre-stimulus. Though, the detection response *×* modality interaction of the dITPC model was not significant at CP4 (*p*_FDR_ = .20), while the main effect of detection response was significant (*p*_FDR_ = .041), indicating that there is no evidence for a different detection effect in the modalities. One reason for weaker phase locking in mechanical stimulation might be that the piezoelectric stimulation timing is less precise compared to the sharp square waveform from the electrical stimulator, which could make exact phase tuning more difficult. Furthermore, the slight shift in timing of the visual cue by 36 ms in the mechanical modality might pose a limitation to the interpretation. However, given the small difference, especially with providing slightly more time in the mechanical condition, we do not believe that it affects the mechanism of perceptual preparation through temporally planned phase locking.

Previous work showed that alpha phase at stimulus onset predicts somatosensory perception [18] and that early post-stimulus phase locking is associated with successful detection [19]. Unlike previous studies employing temporally unpredictable stimulation, our paradigm allows the observed phase alignment to be related directly to the presentation of the preparatory cue. Similar cue-induced phase alignment has been reported in the visual domain, where pre-stimulus alpha phase synchronization is associated with enhanced post-stimulus processing [24]. Our findings suggest that a comparable mechanism may also operate in the somatosensory system when temporal expectations are established by a visual cue, but this requires further testing.

In our study, plotting the P50 amplitude by alpha phase bin, we observed that the highest amplitude was at phase bin 180-202.5° for electrical and 135-157.5° for mechanical stimulation (see Fig. 6 B). Extrapolating this angle to stimulation onset assuming a 10 Hz wave, the oscillation would reach the peak at 47 ms for electrical and 59 ms for mechanical stimulation (for demonstration of the extrapolation see Fig. 6 D). This matches well with the expected time points of the P50 component peak respectively. The pre-stimulus alpha phase at 100 ms before stimulus onset revealed an average phase of 248° in the electrical condition, which corresponds only weakly to the centre of the strongest phase bin (180-202.5°), since it is about 57° apart. In the mechanical condition the alpha phase was not specific to a certain phase at 100 ms pre-stimulus. It is therefore not directly clear if tuning specifically to the P50 peak might enhance perception. Moreover, it is unlikely that pre-stimulus alpha and the P50 share the same neural mechanisms, since their topographies differed significantly (see Fig. 6 C). The effect that pre-stimulus alpha had on stimulus detection had a somatosensory hotspot (electrodes CP4 and C4 in the electrical and mechanical condition respectively), but did not show the same pattern of the perceptual effect at the P50 time point. We speculate that pre-stimulus alpha is a separate process from the generation of the P50 component, but that it might use shared pathways to modulate perceptual processing. The relationship between pre-stimulus alpha and the P50 amplitude is highly complex and depends on attention [3]. In source space, the cluster-based permutation test of the pre-stimulus alpha dITPC effect did not reveal a significant cluster in either modality, so the cortical origin of the sensor-level effect at CP4 remains undetermined. We replicated the effect of alpha amplitude on somatosensory near-threshold detection at the electrode C4, without a significant interaction between modalities [3, 18]. The inverted-U relationship was significantly negative at C4 in the electrical condition (the relationship pointed in the same direction for the mechanical modality but only survived FDR correction at the electrode O2). Though, the curve is mostly monotonic and does not show a high quadratic coefficient. In both modalities, the hotspot of the detection-suppressing effect of the alpha envelope was located in the somatosensory region (CP4 for the electrical and C4 for the mechanical modality), indicating that this reflects Rolandic alpha (Fig. 4C).

Our cluster-based permutation analysis of broadband data revealed a preparation effect that was not specific to the somatosensory modality and was most prominent over central and frontal electrodes. This effect closely resembled the well-studied contingent negative variation (CNV), reflecting anticipation and expectancy [47–49]. The CNV does not guarantee conscious stimulus perception but instead reflects anticipation and attentiveness; it has also been reported in placebo experiments in which participants anticipated pain [50, 51]. In previous studies the pre-stimulus interval was typically longer (mostly exceeding 1 s), yet we nevertheless observed the negativity component under our shorter interval.

In comparison to the pre-stimulus alpha effect, our negativity component differed in two key aspects: it emerged later (from approximately -0.15 s) and at a lower frequency (*<*6 Hz) (Fig. 7A). Furthermore, the negative peak was present in both modalities, whereas the dITPC effect was observed only for electrical stimulation and therefore seems to stem from a different preparatory process. Prior work in other sensory modalities indicates that CNV and alpha-band mechanisms co-occur during anticipation, though this association is typically attributed to alpha event-related desynchronization (ERD) rather than pre-stimulus alpha phase locking [52–54]. Filipovic et al. [54] argued against treating alpha ERD and the CNV as two measures of a single underlying preparatory process. The topography of the CNV appeared to shift from fronto-central to parieto-central electrodes, consistent with observations in the auditory domain [52]. We computed the correlation between the peak pre-stimulus dITPC and the peak of the negativity component for perceived and unperceived trials, by modality. None of the correlations was significant; the data therefore do not show a relationship between alpha phase locking and the amplitude of the negativity in either modality, which is in line with the view that the two reflect different preparatory processes.

## Conclusion

In this study we showed that increased pre-stimulus alpha phase locking is associated with increased stimulus detection. Furthermore, we confirm previous findings that a lower pre-stimulus alpha amplitude goes along with a higher detection rate. There was no evidence that the effects differed significantly between modalities. Post-stimulus neural correlates of conscious perception were identified at the N80 and the N140 component, which appeared largely similar for mechanical and electrical somatosensory near-threshold stimulation. Exploratory analyses suggest that the P50 component amplitude is weakly related to pre-stimulus alpha oscillations in a way that the P50 was enhanced when the alpha oscillation was aligned with its peak. The two originate from different processes, since prestimulus alpha envelope and P50 topographies differed significantly. Future research should further investigate the interaction between alpha-band activity and the P50 component during perceptual preparation and determine how the timing of the preparatory cue shapes pre-stimulus phase locking in the alpha band.

## CRediT authorship contribution statement

**Conceptualization:** RG, SW, TRK, AV; **Data curation:** RG, SW; **Formal analysis:** RG, SW, TRK, AV; **Funding acquisition:** TRK, AV; **Investigation:** RG, SW, TRK, AV; **Methodology:** RG, SW, TRK, AV; **Project administration:** RG, SW, TRK, AV; **Resources:** TRK, AV; **Software:** RG, SW; **Supervision:** TRK, AV; **Writing – original draft:** RG, TRK, AV; **Writing – review & editing:** RG, SW, TRK, AV.

## Declaration of competing interest

The authors have nothing to declare. There are no conflicts of interest.

## Data and code availability

The analysis code is available on GitHub at https://github.com/rosagross/somatosensory_ntdetect. The data are available on Zenodo at 10.5281/zenodo.24078965.

## Appendix

Elimination of visual evoked response applying catch trial subtraction

**Figure 8:**
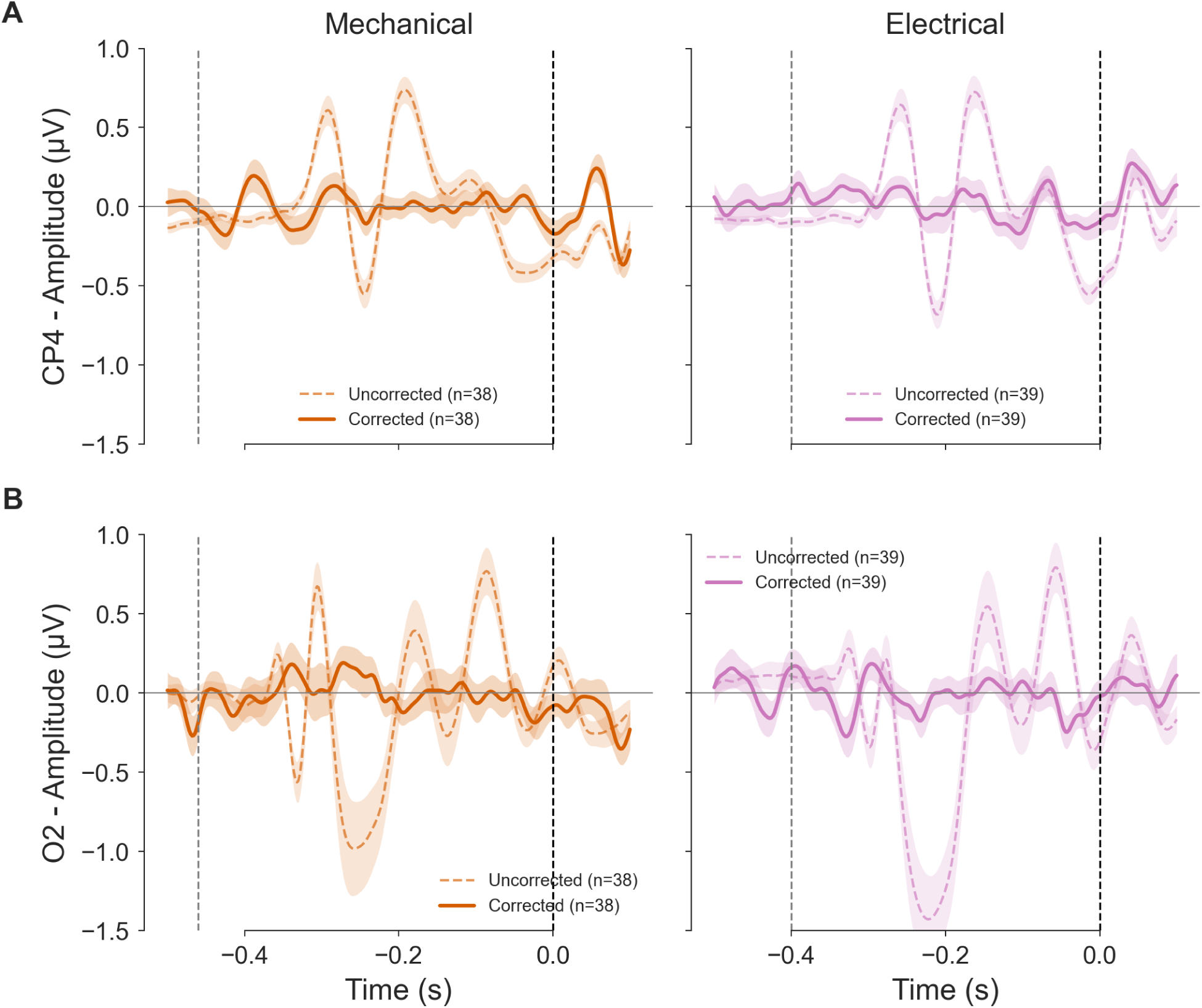
Elimination of visual evoked response applying catch trial subtraction.

## Notes

https://zenodo.org/records/24078965

### Competing Interest Statement

The authors have declared no competing interest.

